# REVEALING COMPLEX ECOLOGICAL DYNAMICS VIA SYMBOLIC REGRESSION

**DOI:** 10.1101/074617

**Authors:** Yize Chen, Marco Tulio Angulo, Yang-Yu Liu

## Abstract

Complex ecosystems, from food webs to our gut microbiota, are essential to human life. Understanding the dynamics of those ecosystems can help us better maintain or control them. Yet, reverse-engineering complex ecosystems (i.e., extracting their dynamic models) directly from measured temporal data has not been very successful so far. Here we propose to close this gap via symbolic regression. We validate our method using both synthetic and real data. We firstly show this method allows reverse engineering two-species ecosystems, inferring both the structure and the parameters of ordinary differential equation models that reveal the mechanisms behind the system dynamics. We find that as the size of the ecosystem increases or the complexity of the inter-species interactions grow, using a dictionary of known functional responses (either previously reported or reverse-engineered from small ecosystems using symbolic regression) opens the door to correctly reverse-engineer large ecosystems.

## 1. INTRODUCTION

Understanding the dynamics of complex ecosystems, such as food webs or human microbiota, has the potential to transform how we approach some of the most pressing challenges of our time, from better ecosystem management to improving human health [1–6]. The human microbiota, for example, is a large and complex community of microbial species primarily residing in the gastrointestinal (GI) tract [7]. Many GI diseases such as *C. difficile* infection, inflammatory bowel disease, irritable bowel syndrome, and chronic constipation, as well as a variety of non-GI disorders as divergent as autism and obesity, have been associated with disrupted microbiota [8–13]. Yet, despite the growing importance of research on those complex ecosystems, there is a remarkable lack of mechanistic understanding of their dynamic behavior. Our uncertainty about the dynamics of complex ecosystems originates in the intrinsic difficulty of extracting useful dynamic models from poorly informative time-series data that we often have. Existing approaches either (i) use parameter identification methods such as multivariate regression [14], maximum likelihood [15] or downhill simplex [16]; or (ii) use a “black-box” framework such as neural or Bayesian networks. In the first case, we must apriori choose the model structure —an assumption that is always hard to justify given the existence of different functional response models[17]. Indeed, this forces us to rely on “standard” models such as the Generalized Lotka-Volterra(GLV) model [4, 18], despite we know its limitations occur even at the scale of two-species [17, 19]. In the second case, despite those “black-box” approaches can offer very accurate prediction of the system’s temporal behavior, they cannot provide any mechanistic understanding of the underlying ecological dynamics.

Here we propose to fill this gap by combining Symbolic Regression (SR) with prior knowledge of possible interaction types (i.e., so-called “functional responses” [17]). As a recent system identification method based on evolutionary computation, SR searches in the space of mathematical expressions both the structure and parameters of an ordinary differential equation (ODE) model that accurately explains the given time-series data [20, 21]. Importantly, SR also provides several candidate models with different levels of complexity and accuracy, letting us choose the model with the best and most significative tradeoff. We show that SR allows us to discover the dynamics of two-species ecosystems with diverse functional responses from time-series data without any prior knowledge of their dynamics, producing dynamic models that can be mechanistically interpretable. Yet, in order to correctly discover the dynamics behind given time time-series data, we find it is essential to have informative enough data. Otherwise, our approach will infer accurate models with dynamics different to those that generated the data. As the size of the ecosystem grows, we find it becomes harder for the data to be informative enough to reveal the full dynamics of the system. In order to circumvent this problem, we propose to use a “dictionary” of functional responses obtained by either reverse-engineering small systems from informative time-series data or from domain knowledge [17]. We validated this method using both synthetic and real data, showing that it can open the door to mechanistically understand the dynamics of complex ecosystems. A schematic overview of the Symbolic Regression workflow discovering the dynamics of a two-species ecosystem is shown in Fig. 1.

**FIGURE 1.**
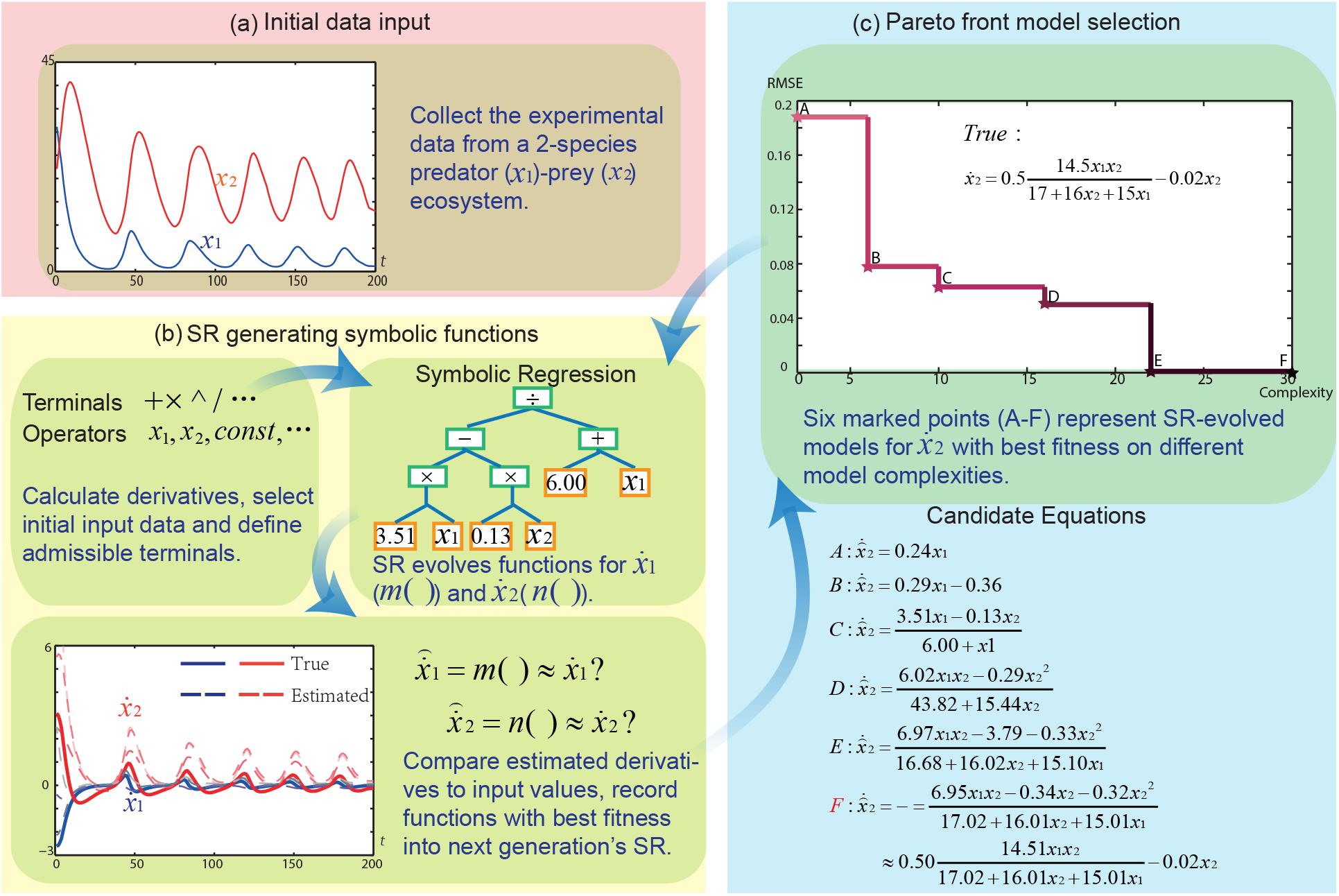
The schematic overview of the Symbolic Regression workflow. **a.** Without any prior information on model structures or parameters, our aim is to find mechanistic understanding of ecological systems given the time-series data input. **b.** The Symbolic Regression algorithm searches a set of functions illustrating the dynamics of the given data, and we use root-mean-square errors (RMSE) to evaluate model fitness. Less RMSE represents higher model fitness. **c.** The Pareto front can reflect the tradeoff between complexity and fitness of candidate equations. With recorded optimal fitness on ceratin value of model complexity, it is meaningful to keep an account of each cliff in the plot corresponding with equation *A,B,C,D,E* and *F*, indicating the increase of predictive ability as model structures evolve. After searching on a space of 1.9 × 10^8^ equations, SR finds equation *F* revealing true model dynamics.

## 2. RESULTS

### 2.1. Two-species ecosystems

Consider synthetic time-series data {*x*_1_(*t*), *x*_2_(*t*)},*t* ∈ [0, *t*_*f*_], generated from a general two-species predator-prey model

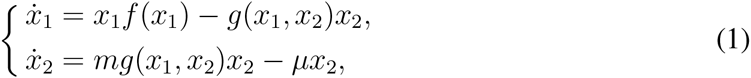

where *x*_1_ and *x*_2_ denote the density of prey and predators, respectively [17]. The function 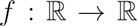 represents the prey growth rate, and 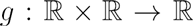 is the so-called “functional response” which describes the instantaneous, per capita feeding rate of the predator and represents the form of interaction between species [22]. The constants *m* > 0 and *µ* > 0 are the conversion efficiency and the per capita death rate of predators, respectively. The standard model for growth rate is given by the logistic equation

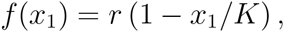

where the carrying capacity *K* > 0 is the maximum number of prey allowed by limited resource, and *r* > 0 is the growth rate constant [17]. Empirical evidence has shown that ecosystems may exhibit very different functional responses [17, 23-28]. Here we consider four representative ones: Lotka-Volterra (LV), Holling Type II (H), DeAngelis-Beddington (DB) and Crowley-Martin (CM):

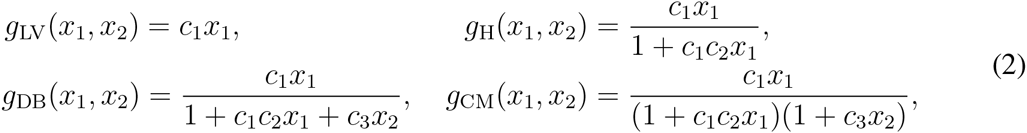

where *c*_*i*_ > 0 are constants. These functional responses describe different *mechanisms* for the inter-species interactions with increasing complexity, which are key factors in determining ecological dynamics (Remark 1 in SI-2.1).

We generated synthetic time-series data by numerically integrating (1) using different functional responses in (2). Then, we used SR to reconstruct 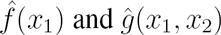 from this data (see Methods), providing estimates for the true *f*(*x*_1_) and *g*(*x*_1_, *x*_2_). The only prior knowledge used in the SR algorithm is that 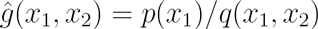 for some unspecified functions *p* and *q,* preventing the SR algorithm from searching over functional responses that are not ecologically meaningful (Methods 3). In order to test the performance of SR, we considered two case studies in which the data have different levels of “informativeness”. In the first case, the parameters *m*, *μ*, *r*, *K* and *c*_*i*_ are chosen such that the systems exhibit a limit cycle (i.e., stable oscillations). In such case, the data was informative enough in the sense that SR was able to correctly recover the functional form as well as parameter values for the LV, H and DB functional responses (Fig. 2a-c). For the CM functional response, SR finds an accurate model (i.e., fits the data accurately), but the inferred functional response 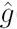 does not match the correct functional response *g* that was used to generate the data (Fig. 2d). This means that the data is still not informative enough to reveal the correct functional response, since different model structures can fit the data equally well. To resolve this problem, more information is needed, and a method often used in practice is to collect time-series data from the response of the prey *x*_*1*_(*t*) in isolation [17]. This extra information allows us to infer *f*(*x*_1_) first, and then to recover *g*(*x*_1_, *x*_2_) (Methods 3). Following this process, the correct functional response can indeed be recovered even in the case of CM interactions (Fig. 3).

**FIGURE 2.**
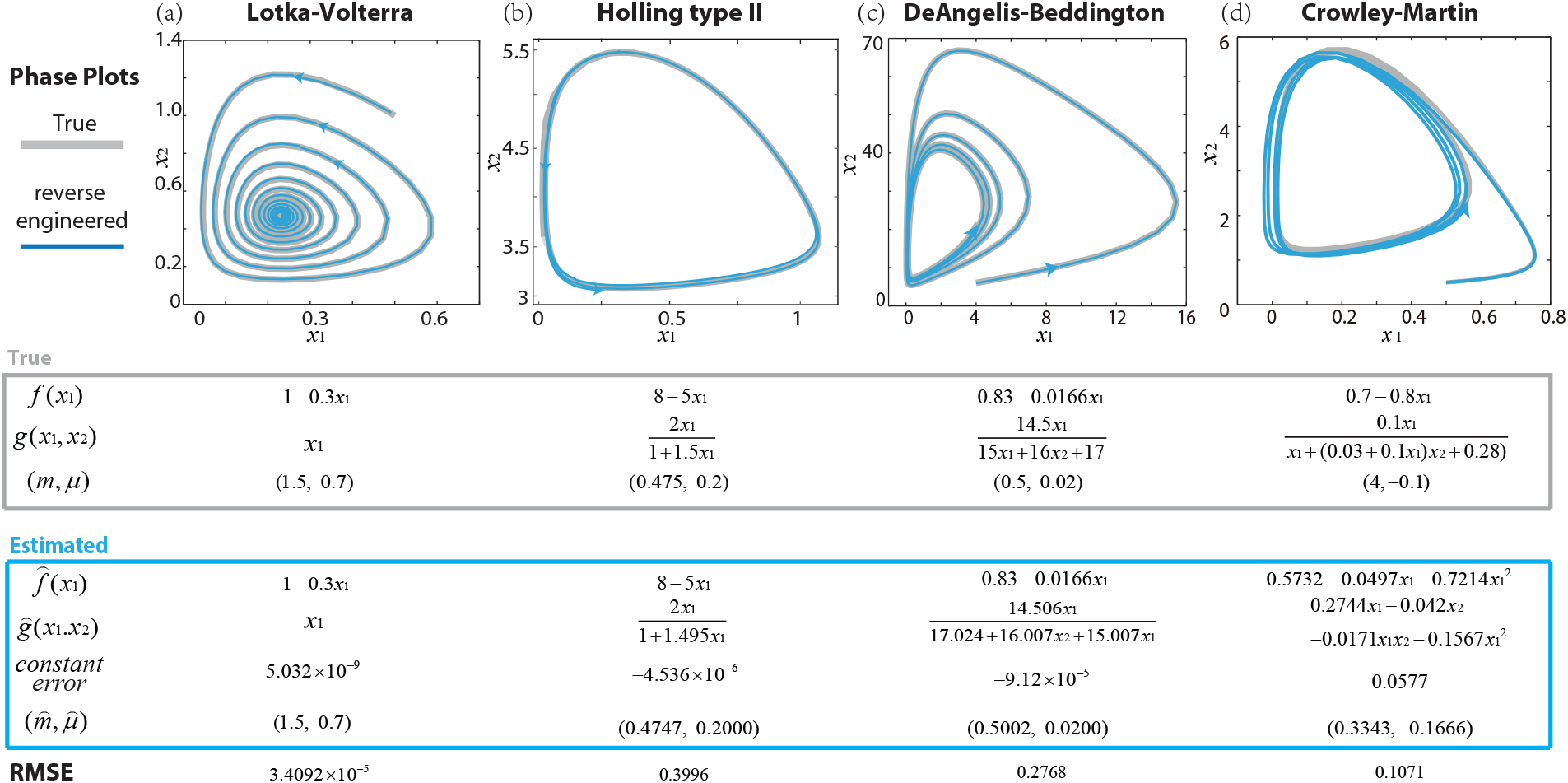
Reverse-engineering synthetic two-species ecosystems. For Lotka-Volterra, Holling type II and DeAngelis-Beddington functional responses with limit cycles, SR can directly reconstruct the correct growth functions and functional responses from time-series data. For Crowley-Martin type with more complex functional response, SR only reconstructs a model with high accuracy but incorrect model structure. Root-mean-square errors (RMSE) are calculated to compare derived models with original synthetic ones, while constant errors are the constant terms in the derived SR models.

**FIGURE 3.**
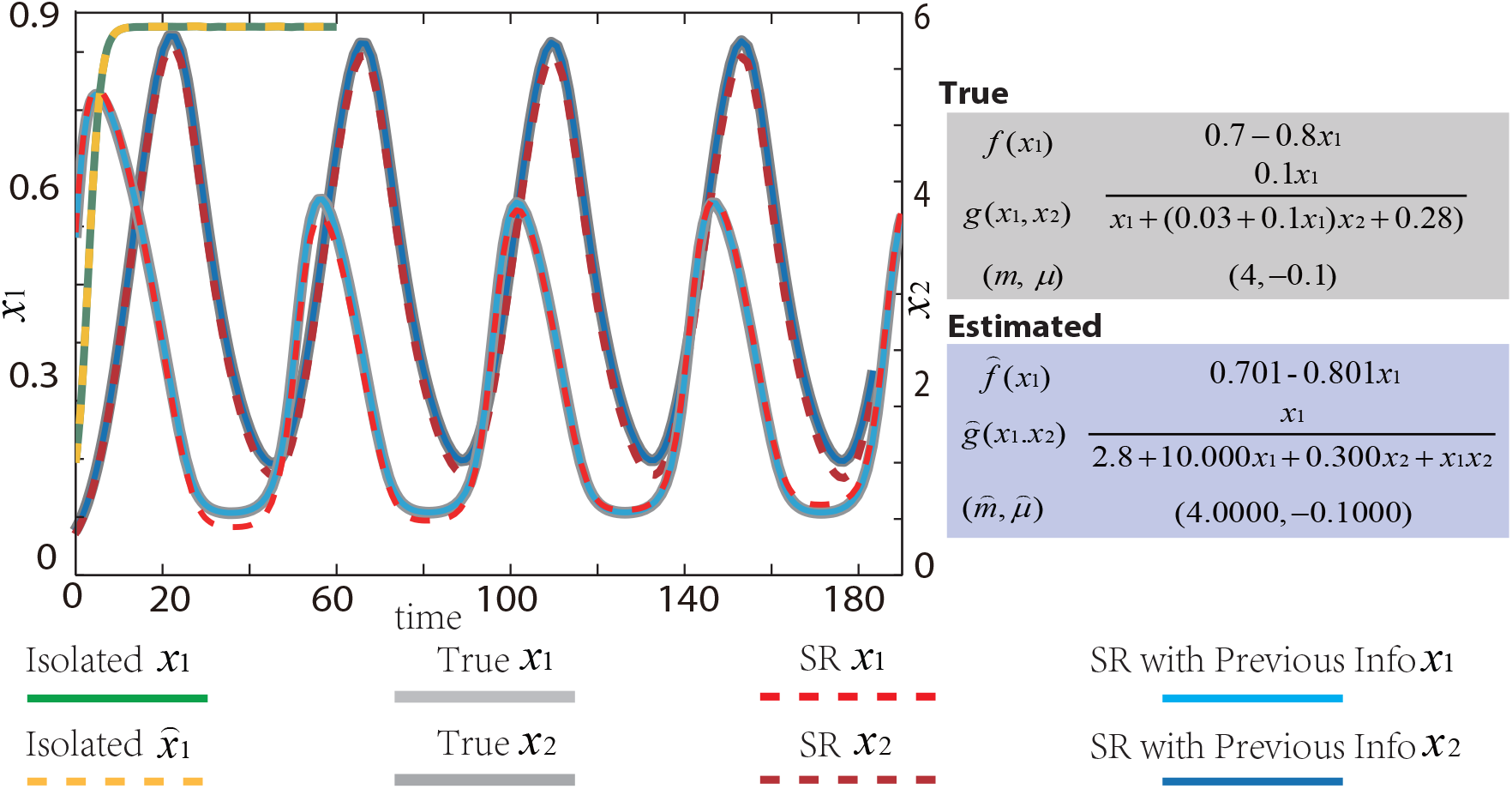
Reverse-engineering a two-specie ecosystem with Crowley-Martin functional response. Without giving any prior knowledge of the interaction (red), SR is able to infer an accurate model that does not use the CM functional response. In order to reverse-engineer the correct functional response, we provide extra information using the time-series data of the isolated prey (green) that allows us to correctly infer *f*(*x*_1_) first (yellow). With such prior information (blue), SR correctly recover the functional form for *g*(*x*_1_,*x*_2_).

In order to better study the role of the informativeness of the measured temporal data on the discovered dynamics, in the second case study we choose the parameters of the system such that its trajectories approach an equilibrium, Fig. 4. The synthetic data obtained in such a way has no persistent oscillations, and SR finds accurate models but their growth rates and functional responses differ from the true ones (solid blue lines in Fig. 4). To circumvent this fundamental limitation, in Section 2.2 we show that prior knowledge of the functional form of the interactions is extremely useful, letting us recover the correct dynamics from otherwise uninformative time-series data.

**FIGURE 4.**
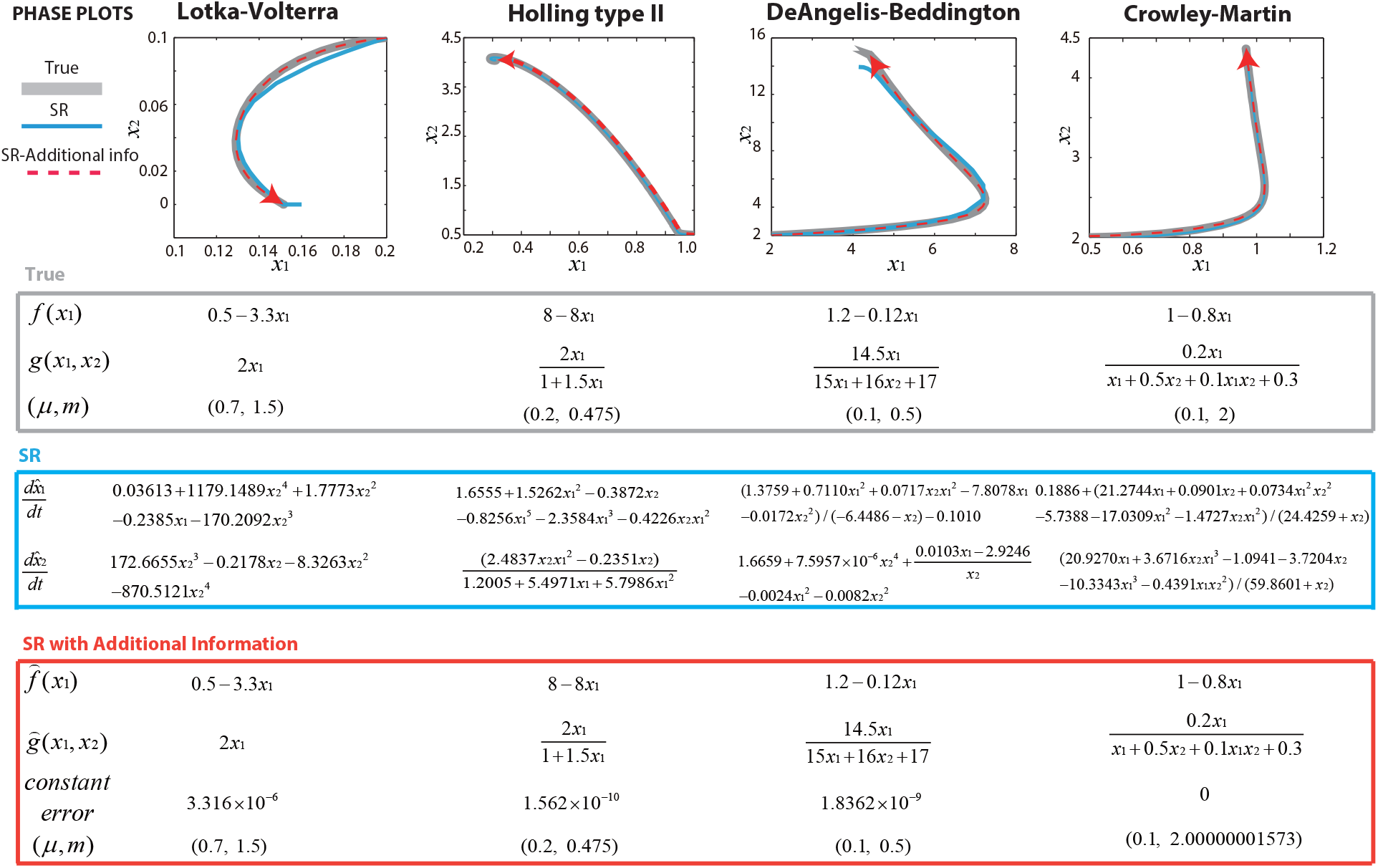
Reverse-engineering a two-species ecosystem from uninformative data. Compared to Fig.2, here the parameters of the system are such that its trajectories quickly approach an equilibrium. From this data, SR is able to reverse-engineer an accurate model without recovering the correct functional response or growth rates (blue). In this sense, the data itself is not informative enough. In order to acquire more information without needing more data, we provide to the SR algorithm a “dictionary” of the possible functional responses. With this additional information, the SR algorithm is able to correctly reverse-engineer both the growth rate and functional response (red).

Next we test our approach with real data from a predator *(P.aurelia)* and prey *(D.nasutum)* ecosystem [29]. Following the methodology of [17], we first infer the growth rate function 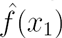 from experimental data of the prey growing in isolation, and we let 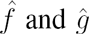 depend on delayed values of *x*_1_ and *x*_2_. Using the SR method, we infer the model

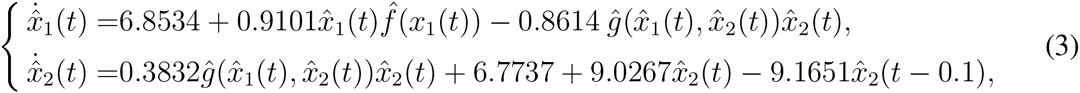

with the following growth rate and functional response

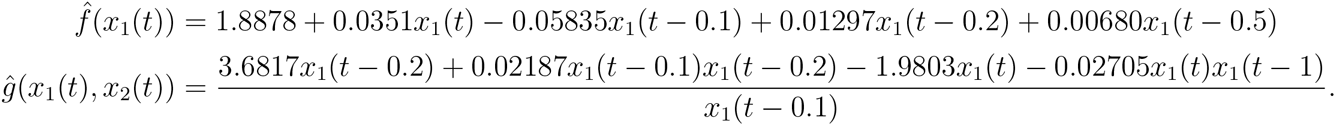

Here the time *t* is in units of days. The inferred model contains constants in the right-hand side of the differential equation for prey and predator, which can be interpreted as external (constant) inputs from the environment acting on the system. The growth rate function 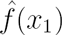 includes several terms with delays in addition to the standard logistic model. For the death rate of predators, the model also includes delays. These delayed terms indicate that the current population affects the carrying capacity of their offsprings. Furthermore, the inferred functional response depends only on the prey. The inferred model (3) using our SR approach has a Root Mean Squared Error (RMSE) of 22.7123, while the best fitted model computed in [17] with DeAngelis-Beddington functional response has an RMSE of 53.4867. Note that such model also contains delays. This means that SR was able to automatically infer a model with more than twice the accuracy, as can be also appreciated by visual inspection of the true and predicted trajectories (Fig. 5).

**FIGURE 5.**
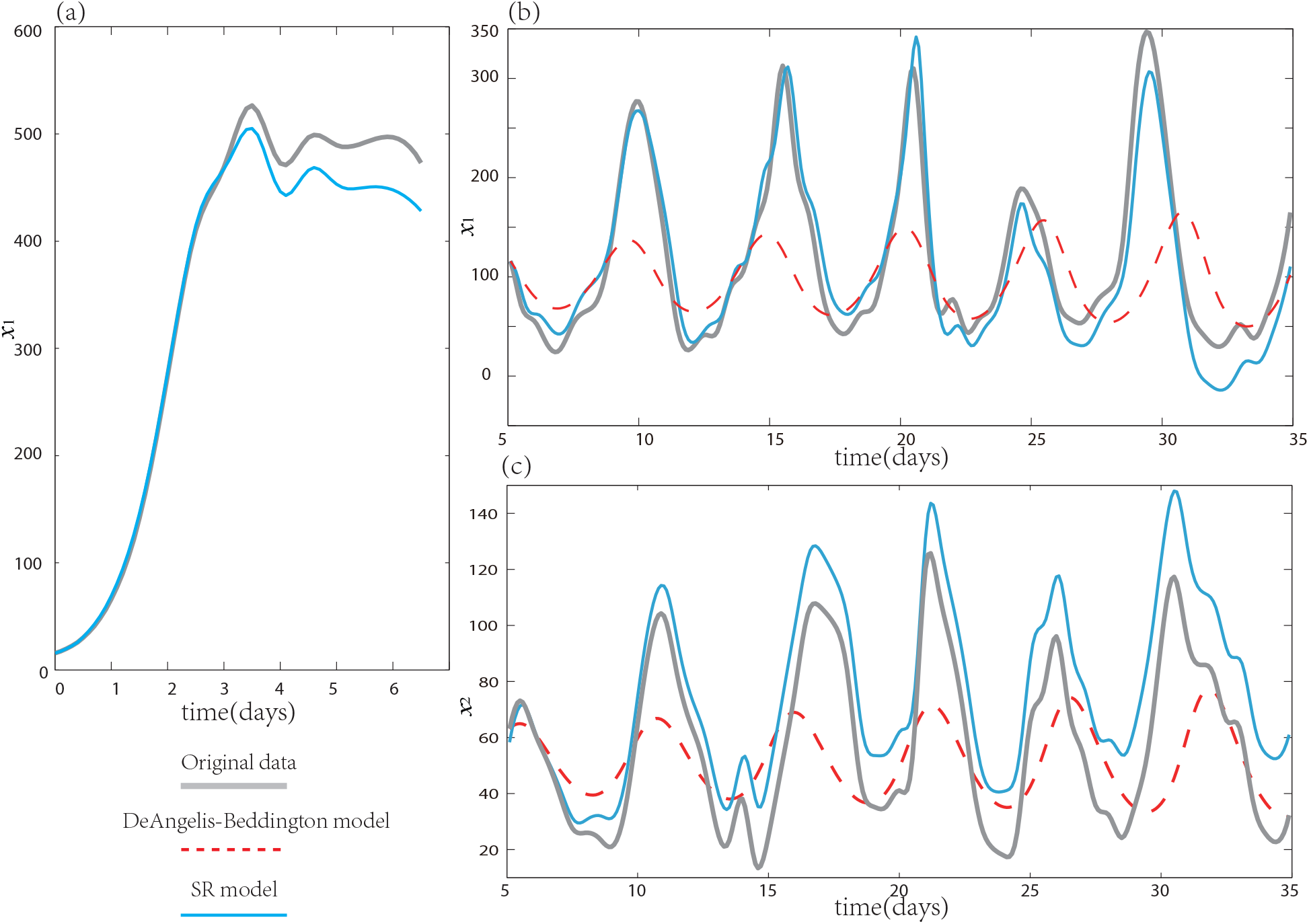
Reverse engineering a predator-prey ecosystem from experimental time-series data. **a.** Experimental time-series obtained from the prey in isolation (grey), and the estimated time-series from the reverse-engineered model using SR (blue). **b.** Experimental time-series data of the prey. True (gray), reverse-engineered model using SR (blue), best model fitted in [17] using the DeAngelis-Beddington functional response with logistic growth (dashed red). **c.** Time-series data of the predator. True (grey), reverse-engineered model using SR (blue), best model fitted in [17] using the DeAngelis-Beddington functional response with logistic growth (dashed red).

### 2.2. Using prior knowledge of functional form of interactions

In the simulation examples of the previous section, we found that if the data is not informative enough then SR can reverse-engineer an accurate model in terms of trajectory prediction, but the model itself is totally different from the ground truth that was used to generate the synthetic data. In order to circumvent this limitation and recover the correct functional response and growth rate, we propose to seed the SR algorithm with a “dictionary” of possible functional responses, Methods 3. This dictionary is built from either previously reported or reverse-engineered from informative enough data using SR. With this additional information, SR can correctly reverse-engineer the correct functional response even with the less informative data of Case 2 in Section 2.1, Fig. 4. Indeed, this prior information is instrumental to infer the dynamics of larger ecosystems because, as the size of the ecosystem grows, it becomes harder for the data to be informative enough to reveal the full dynamics of the system.

### 2.3. Reverse-engineering larger ecosystems

Finally we test our framework in larger ecosystems, generating data by simulating the following model with six species:

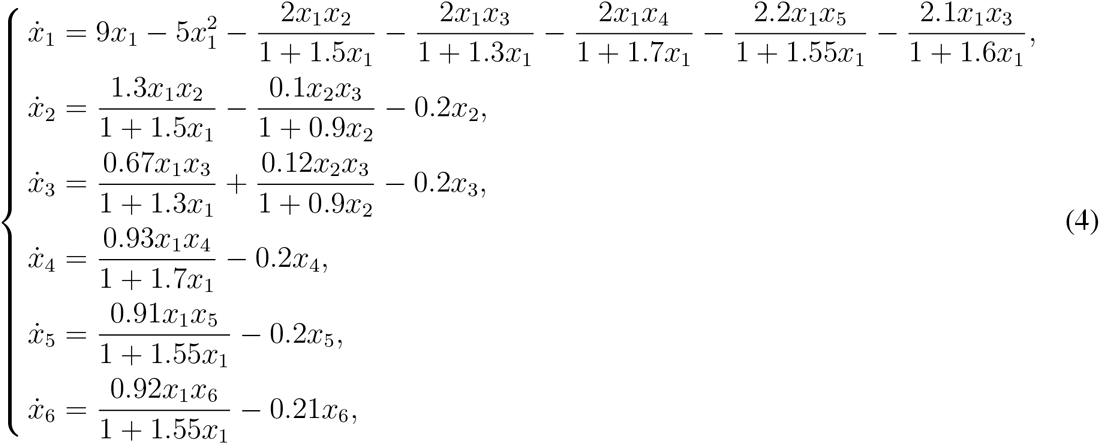

whose interactions are of Holling Type II. We selected the parameters of this system such that its trajectories oscillate as shown in Fig. 6a. By applying SR directly, we obtain an accurate model but it does not contain the correct form of the interactions, Fig.6c and SI-4. Indeed, from Fig. 6a, the time-series of the variables *x*_4_, *x*_5_ and *x*_6_ are very similar, making difficult for any algorithm to distinguish between them (in other words, the effect of including any of these variables in the right-hand side of an ODE model is very similar). Furthermore, SR often yields accurate but very complex models (Methods 3 and dashed line in Fig. 6b). These problems are circumvented by using the dictionary of possible functional responses described in Results 2.2. With this prior information, SR is able to significantly decrease its searching space, and reverse-engineer an accurate model with the correct interactions (Methods 3 and solid line in Fig. 6b).

**FIGURE 6.**
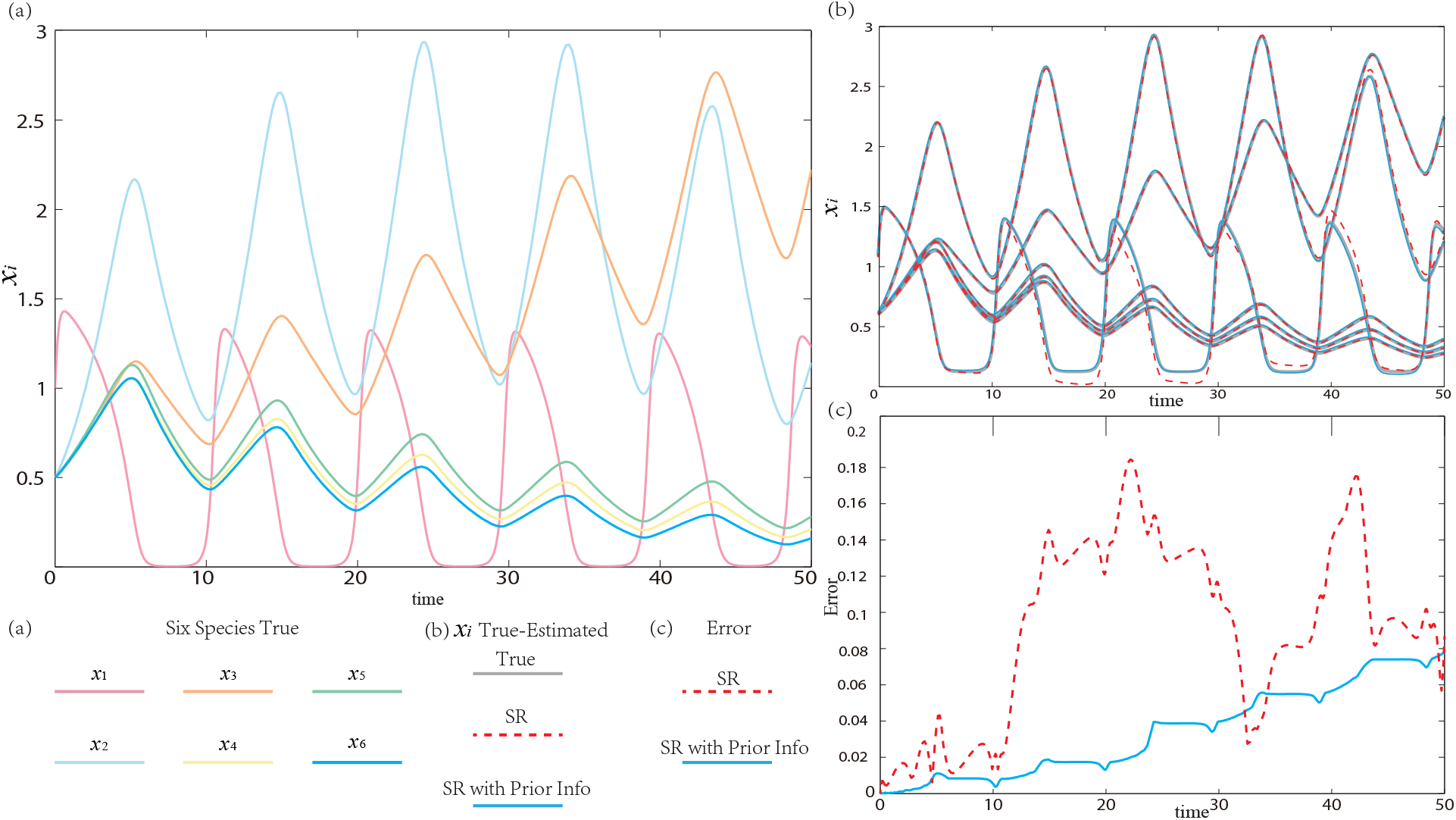
Reverse-engineering larger ecosystems using symbolic regression. **a.** The data consists of the trajectories obtained by simulating system (4) containing six species (each species shown in different color). **b.** True trajectories (grey), trajectories estimated by the reverse-engineered models using symbolic regression without (dashed) and with (solid) prior information. **c.** Error (euclidean norm of the difference between the true trajectory *x*(*t*) and the estimated trajectory 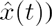 as a function of time for the reverse-engineered models using symbolic regression without (dashed) and with (solid) prior information of the possible interaction types. In both cases, the reverse-engineered models have good accuracy but only when the SR is given prior information the data is informative enough to recover the correct functional responses.

## 3. DISCUSSION AND CONCLUDING REMARKS

There is an increasing need to understand the dynamics of complex ecosystems. Here we introduced a novel method based on SR that is able to reverse-engineer ODE models from time-series data of ecological systems. In particular, with sufficiently informative data, our approach can recover both the structure and the parameters of a model that accurately explains the data. This performance is not shared by most system identification algorithms since, even if the data is informative enough, they can at best fit the parameters of an a-priori selected model (the selection of such model is hard to justify in practice) and often, even if the model accurately explains the data, such models do not provide mechanistic understanding of the system (e.g., in neural network models). Moreover, the proposed SR approach has an additional degree of freedom: it lets the user choose the model that has the best tradeoff between complexity and accuracy for each particular application.

With uninformative data, our approach discovers different ODE models (with different functional response and growth rate functions) that explain the data equally well. This implies that the “true” system dynamics is unidentifiable from the given data [30], reflecting a fundamental limitation to infer the correct dynamics using any method. We found that the informativeness of the data decreases with the complexity of the interactions between species and with the number of species. In order to increase the informativeness we can acquire more data (e.g., time-series from the prey in isolation) and use a dictionary of prior information of the possible functional form of the interactions (functional responses). By seeding the SR algorithm with a dictionary of possible functional responses, we found it is possible to correctly reverse-engineer more complex and larger ecosystems for which the data alone is not informative enough. In particular, in the case of experimental data, we found our approach can produce models twice as accurate as the best model previously fitted.

## METHODS

### Reverse-engineering dynamic systems using symbolic regression

Consider we are given time-series data 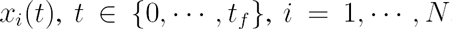, for the abundance of each of the *N* species composing the ecosystem. Our objective is to find functions 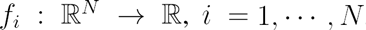, such that the model

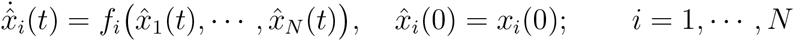

accurately explains the mechanisms behind the data: 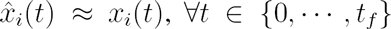 and 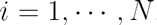.

We are particularly interested in functions *f*_*i*_ that simultaneously are (i) simple (i.e., they can be constructed with the least number of operations), (ii) meaningful from an ecological perspective, and (iii) have good fitness. Here the fitness of a given function *f*_*i*_ is defined using the root-mean-square error (RMSE) between the true and estimated derivatives

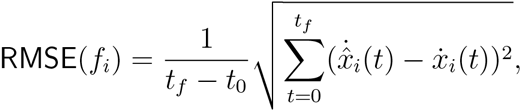

where 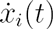 is the estimated derivative of the time-series data and 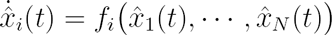. A good tradeoff between these three characteristics yields simple and powerful models, which can be interpreted to understand the dynamic behavior of the ecosystem. SR starts by randomly assembling several candidate a function *f*_i_ using the set of admissible operators {+, −, ×} and terminals {*x*_1_,…,*x*_*N*_} ∪ {const.}. Next, the SR algorithm computes the fitness of each candidate function, keeps the better ones, and uses mutation and crossover [31] among these functions to build better ones [21] with evolution in structures and parameters. This process is iteratively repeated until sufficiently “good” functions are found. In order to achieve this, it is very useful to keep track of the so-called Pareto front that plots several models according to its complexity and fitness, see Fig. 1c.

Unlike typical regression methods like second-order polynomials that specify a model structure with model’s parameters adjusted to fit the data, SR can infer both the model structures and the parameters. In particular, since the functions *f*_*i*_ in ecological models tends to be the sum of small nonlinear functions (i.e., sum of functional responses for each species), multi-gene algorithms [32] are useful.

Expressing models in multi-gene approach uses several genes combined together to evolve equations containing many variables, and it also carries benefits for analyzing the Pareto front, since we can clearly record improvements in the accuracy and complexity of the functions [33]. With such Pareto-aware SR algorithms, we can explicitly explore the trade-off between model complexity and accuracy, letting us select those models that provide the best balance between accuracy and complexity.

### Applying SR to ecological systems with two species

Given time-series data of the two species {*x*_1_(t), *x*_2_(t)}, we first estimate their derivatives 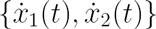 using central difference method. Next, we reverse-engineer a model that accurately fit 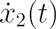. For this, we use SR to find a function 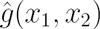 such that

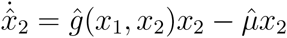

has good fitness/complexity tradeoff for some constant 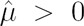. Finally, with the function 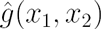, we use SR again to find a function 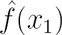 such that

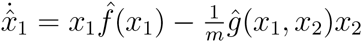

has again a good fitness/complexity tradeoff for some constant *m* > 0. For the results shown in all figures, we used the SR algorithms incorporated in Eureqa [34]. Eureqa also lets us incorporate the constraint 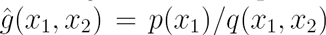 for some function *p* and *q,* preventing the SR algorithm to search over model spaces of functional responses that are not ecologically meaningful.

### Prior information I: additional data from the prey in isolation

We explored two methods to incorporate prior information. The first one uses more data from the response of the prey *x*_1,isolated_ (*t*) in isolation. We estimate again the derivative of this data 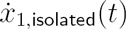 and use SR to find a good function 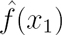 such that

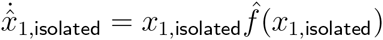

has a good fitness/complexity tradeoff. With this estimated 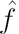, we can use the time-series data from the prey interacting with the predator {*x*_1_(*t*), *x*_2_(*t*)} to reverse-engineer the functional response 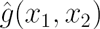 by using

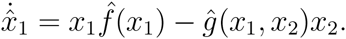

We found this approach very efficient, reducing the time required to correctly reverse-engineer 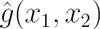, which in turn reduces the reverse-engineering process to find parameters *m* and *µ.* This method allows us to correctly reverse-engineer a synthetic ecosystems of CM functional responses (results shown in Fig. 3b).

Since we can expect that the functional response of real ecosystems are at least as complex as the CM functional response, we applied the same method to the experimental data of Veilleux [29]. We exploit interpolation and delay operator to build the candidate functions for the SR algorithm.

### Prior information II: prior knowledge of possible functional responses

The second method to incorporate prior information simply seeds the SR algorithm with prior knowledge of possible functional responses. Instead of trying to reverse-engineer the equation for 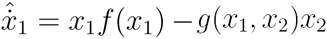, we listed all possible units which may exist in the denominator of *g*(*x*_1_,*x*_2_), like *ax*_1_, *bx*_2_, *cx*_1_*x*_2_ and *dx*_2_, and treated them as a dictionary of interaction forms for inputs of SR. In the next step, we transformed the reverse-engineering process of 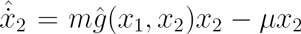 into a multi-gene SR problem of finding parameters for different units in our interaction dictionary. Some parameters simply equal to 0, indicating the non-existence of some types of functional responses. We performed the multi-gene symbolic regression using the Matlab package GPTIPS2 [35], combined with a post-analysis on Pareto-front to select the best transformed model with the fintness/complexity tradeoff. From a technical perspective, compared to previous symbolic regression procedures, we found that combining the dictionary of possible interactions with multi-gene genetic programming increases the accuracy of the method and helps avoid bloated equations (i.e., accurate but extremely large models). Nevertheless, this choice tends to produce models with a small constant error that is accumulated when the ODE models are integrated. This could be remediated by using a different norm for evaluating the fitness of the candidate models in the SR algorithm. Such choice, however, would slow down the SR algorithms because it requires to numerically integrate an ODE system to evaluate the fitness of a candidate model.

This approach is pretty useful in selecting the true functional response for uninformative data and finding the model for 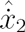, and then we follow the same step in Methods 3 to reverse-engineer 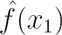. Thus prior information on the possible functional responses proves to be very useful in recovering the model structures, especially for those uninformative data sets.

### Applying SR for larger ecosystems

We comprehensively employed prior information mentioned in Methods 3 and Methods 3. In our results shown in Fig. 6, at the initial stage, as *x*_4_, *x*_5_ and *x*_6_ all conforms to the model structure of

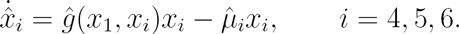

SR can directly reverse-engineer 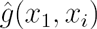 and *µ*_*i*_ by only providing time-series data *x*_1_(*t*) − *x*_6_(*t*) as inputs, which is the same case in recovering 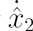 in Methods 3. It selected out the related variables to the derivative 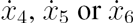 and successfully reverse-engineers the ODEs.

We then found SR was stuck at finding the correct models for the dynamics of *x*_2_, *x*_3_ if we provided no more knowledge of model itself, as it had three species included in one ODE. At this stage we listed all the possible forms of interactions described in Methods 3, and instructed SR to use this dictionary as the prior knowledge for model reconstruction. Then multi-gene SR helped selecting existing forms of functional responses, and reverse-engineering 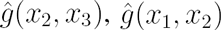 and 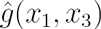. With all the recovered functional responses 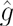 concerned with *x*_1_ provided as inputs, the SR algorithm was able to correctly infer the model of 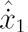, which has the highest complexity including pairwise interactions with other 5 species. The typical three steps of revered engineering plots are shown in Fig. 6b, with a comparison to the original synthetic model and direct SR model with good fitness but poor structures.

## Acknowledgements

This work was supported by the CONACyT postdoctoral grant 207609 and the John Templeton Foundation: Mathematical and Physical Sciences grant no. PFI-777.

## Author Contributions

Y.-Y.L. and M.T.A conceived and designed the project. Y.C. performed all the numerical calculations and data analysis. All authors analysed the results and wrote the manuscript.

## Author Information

The authors declare no competing financial interests. Correspondence and requests for materials should be addressed to Y.-Y.L.

## 1. PRIMER ON SYMBOLIC REGRESSION

Based on genetic programming [1], symbolic regression (SR) is a methodology to search in a space of mathematical expressions for those that accurately fit given temporal data [2, 3]. Note that SR is able to search for both the parameters and the functional form of such expressions, letting us build models based on Ordinary Differential Equations (ODEs) [2] for dynamical systems. In order to perform SR, we need to predefine a set of admissible operators (binary operations like {+, −, x } and unary operations like {*log, exp*})and the set of “terminals” 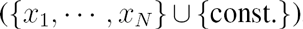, which the algorithm can use to build mathematical expressions. For example, the function *f*_*i*_(*x*_1_,…,*x*_*n*_) = 2*x*_1_ + 1.6 requires two operators and three terminals.

In the initial stage, the classical SR algorithm randomly generates assigned number of candidate functions {*f*_*i*_} combining randomly a subset of terminals and operators. *The fitness* of each of those candidate functions is computed, quantifying how fit the data (see Methods in the main text for details). In addition, model-building information for each evolved equation, such as function complexity and individual fitness are also recorded as criteria in selecting meaningful while concise candidates during the searching process. In the next stage, the SR algorithm keeps the candidate functions with better fitness, and uses evolutionary computation [1] to construct “better” candidate functions from them. This is done via two methods: *mutation* (alters, deletes or adds an terminal or operator to an existing function) and *crossover* (creates two new offspring functions for the new generation by genetically recombining randomly chosen parts of two selected parent functions). This process is iteratively repeated until models with high fitness and low complexity (measured by number of operators and terminals used) are found. Using the “Pareto front” —a plot of inferred models according to their complexity and fitness— SR algorithms are able to efficiently track and control this process.

Note that unlike typical regression methods in which a model structure must be a-priori fixed (e.g., second-order polynomials, wavelets, sigmoids, etc.) and only the model’s parameters are adjusted to fit the data, SR can infer both the structure of the model and its parameters simultaneously. In other words, SR algorithms let us search over the (infinite dimensional) space of possible ODE models for those accurately fitting the data. It has also been shown that incorporating intermediate regression and ensemble steps, such as providing a group of sigmoid functions or selecting the most representative candidates during a generation, can enhance and accelerate its performance [4].

An intrinsic drawback of SR is that the search spaces increases exponentially as the number of terminals or operators increases leading to more complex equations. This implies that it becomes harder for SR algorithms to correctly infer the interactions between species (i.e., functional forms) as the number of species increases or as the interactions become more complex. Nonetheless, since reported functional responses in ecological systems tend to be linear combinations of rather simple nonlinear functions [5], we found that a variant of traditional SR known as multi-gene algorithms [6] can be very useful. Instead of using a single genetic programming tree that easily becomes very large with complicated structure in each of its branches, in multi-gene SR we evolve simultaneously several (independent) trees restricting their complexity. Trees represent genes that can be combined to build candidate equations and hence candidate ODE models. The multi-gene approach is also useful for analyzing the Pareto front, since we can more easily record improvements in the accuracy and complexity of the functions [4]. Indeed, we can decompose the equations on the Pareto fronts during each run, helping us extract sub-blocks (e.g., *x*_*i*_*x*_*j*_ or *x*_*i*_*x*_*j*_/(const. + *x*_*j*_)) that recurrently appear in the interactions between different species. This allow us to explicitly explore the trade-off between model complexity and accuracy by select those models that provide the most useful balance between accuracy and complexity.

## 2. SYMBOLIC REGRESSION TO INFER MATHEMATICAL MODELS OF ECOSYSTEMS

Previous studies have focused on establishing a useful class of mathematical models than can describe ecological systems [7, 8]. A general class of such models can be written as the following set of ODEs

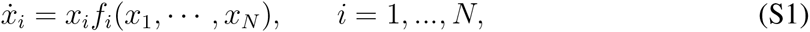

where *x_i_* represents the state (e.g., abundance) of the *i*-th specie in a community of *N* species. The properties of such models provide useful information about the mechanisms behind ecosystems, from stability to the existence of periodic orbits as well as model chaos. Therefore, given temporal data of each species in the system 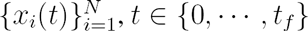, we aim to find functions 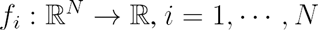, such that the model

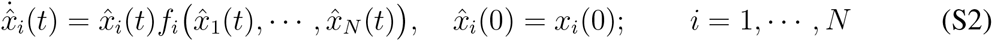

accurately fits the data: 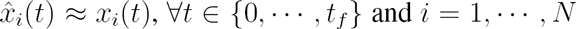. Since, in principle, there is an infinite number of such functions, it is useful to discriminate between them according to their complexity and fitness. In other words, we will be interested only in those functions {*f*_*i*_} which have low complexity and high fitness.

### 2.1. Two-species Ecosystem Dynamics

Consider a general two-species predator-prey model:

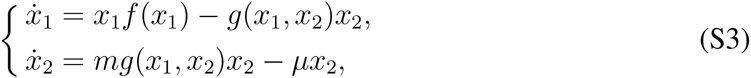

where *x*_1_ and *x*_2_ denote the density of prey and predators, respectively [5]. The function 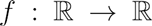 represents the prey growth rate, and 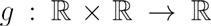 is the so-called “functional response” which describes the instantaneous, per capita feeding rate of the predator and represents the form of interaction between species [7]. The constants *m* > 0 and *µ* > 0 are the conversion efficiency and the per capita death rate of predators. Different forms of functional responses represent different distribution of predators through space as well as the stability of predator-prey systems [9].

The standard model for growth rate is given by the logistic equation

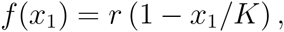

where the carrying capacity *K* > 0 is the maximum number of prey allowed by limited resource, and *r* > 0 is the growth rate constant [5]. Empirical evidence has shown that ecosystems may exhibit very different functional responses [5, 9-13]. Four representative ones are the Lotka-Volterra (LV), Holling Type II (H), DeAngelis-Beddington (DB) and Crowley-Martin (CM) interactions. Their structure is as follows:

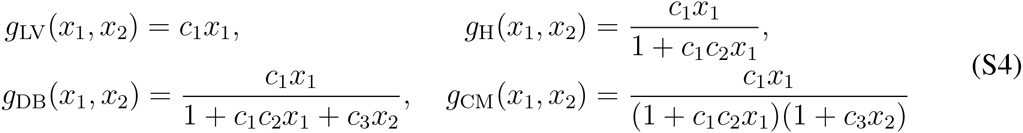

where *c*_*i*_ > 0 are constants. In Fig. S1 we also show the parameters we use in our synthetic models of different types of interactions. Other types of functional responses like Holling Type III, Hassell-Varley [14] and Holling-Tanner [15] have structures similar to these four ones or include less complex interactions. Therefore the successful dynamics discovery of these four functional responses can be fundamental to the research of other ecological models.

**Remark 1.** Different functional responses in (S4) correspond to different *mechanisms* of interaction between species. In the Lotka-Volterra model, the rate of predation is proportional to the rate of instantaneous number of predator. The functional response of Lotka-Volterra model is also regarded as the Holling Type I with the linear increase on food density. On the other hand, in the Holling Type II model, the predator spends time searching and processing the prey. Indeed, the parameter *c*_1_ encodes the effects of capture rate and *c*_1_*c*_2_ describe the effects of handling time for captured prey. In the DeAngelis-Beddington functional response, the parameter *c*_3_ is added to model interference between different predators. The Crowley-Martin models “preemption” allowing for interference among predators regardless of whether a particular individual is currently handling prey or searching for prey. Therefore, by inferring the functional response using SR algorithms, we can learn the ecological mechanisms behind given time-series data.

### 2.2. The role of the informativeness of the data

In order to discover the true dynamics behind given temporal data, it is crucial that the given data itself is informative enough. Otherwise, different dynamics (e.g., models with completely different functional responses) can all fit precisely the same temporal data.

On one hand, when the number of samples is limited, it is usually reluctant to reveal the overall temporal characteristics of each species that decreases data informativeness. On the other hand, we tuned parameters for different types of functional responses based on [16, 17] to produce different time-domain characteristics, and found that informative enough data can be obtained when it records oscillations in the time-series trajectories of the system in the first row of Fig. S1. If the trajectories of the system simply converge to an equilibrium point as shown in the second row of Fig. S1, the data is considered as not informative enough, in the sense that the SR algorithms are able to find ODE models that correctly fit the data yet having different structures of functional responses. For the ODEs we want to recover, such states of quick equilibrium could reflect little of the functional responses’ characteristics itself, while on the other side, from the perception of SR algorithm, it can find a set of eligible candidate ODE models, which fit the original temporal data equally well. Indeed, in such case, we find that it is often possible to fit the data using the simple LV functional response to some extent (Fig. S2 green). This result also helps us explain the wide-spread use of the LV to model diverse ecological systems, because the simple LV model can roughly depict the oscillating and periodic dynamics of ecological systems.

**FIGURE S1:**
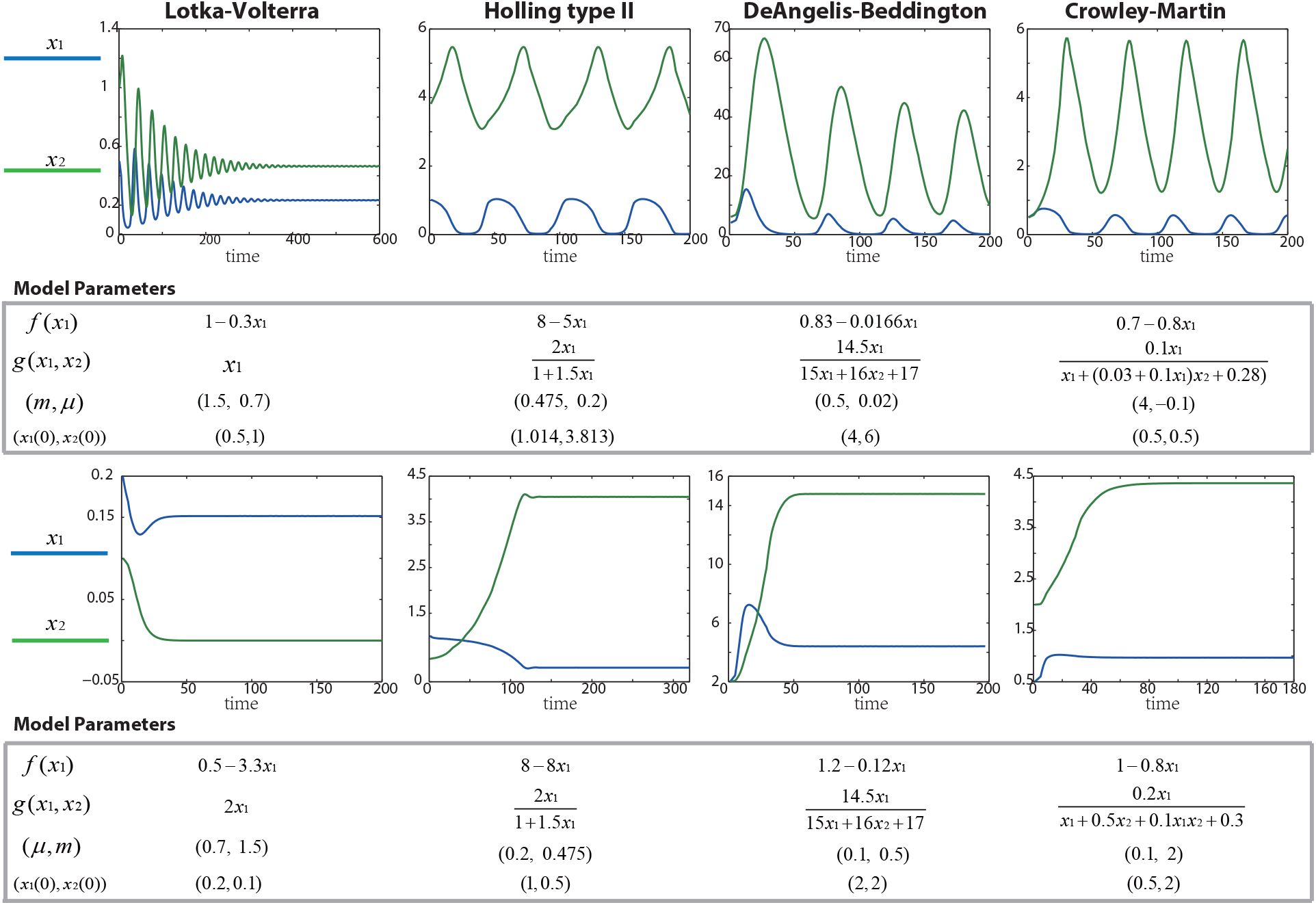
Model inference using informative and uninformative data (same as Fig. 1 and 3 of the main text). Parameters and functional form of the ODE model, and its initial conditions. (a) With oscillations in the trajectories, the temporal data is informative enough for the SR algorithm to recover the correct functional response. (b) Without oscillations, the data is not informative enough and the SR algorithm recovers different functional responses.

**FIGURE S2:**
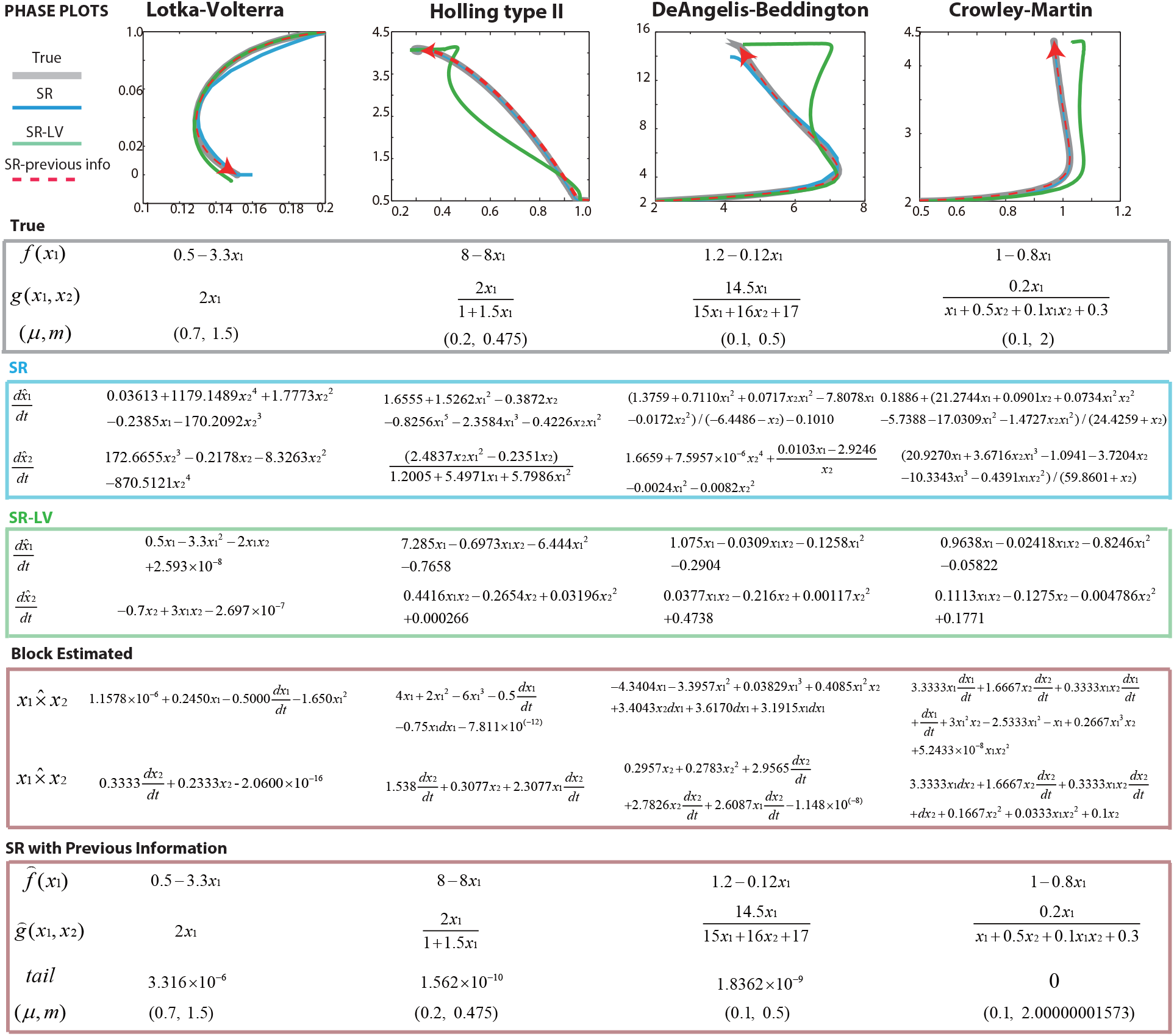
Comparison of direct SR, SR with LV functional response, and SR with a dictionary of functional responses. We consider temporal data in which the trajectories of the system converge to an equilibrium point. The SR algorithms discovers accurate models that have different structures (blue). Indeed, we can force SR to derive models containing the Lotka-Volterra functional response showing the ability of the LV model to fit the response of other functional responses (green). Transforming the model into a linear regression form and using a dictionary of possible functional responses, SR can efficiently infer the growth-rate and functional response even with poorly informative data (red).

Next we move further to build a model for a six-species food web. Based on our analysis of the case of two species, we carefully designed the interactions its parameters in to produce oscillations that could potentially be informative enough to reveal the correct interactions between species. With these considerations, we obtained

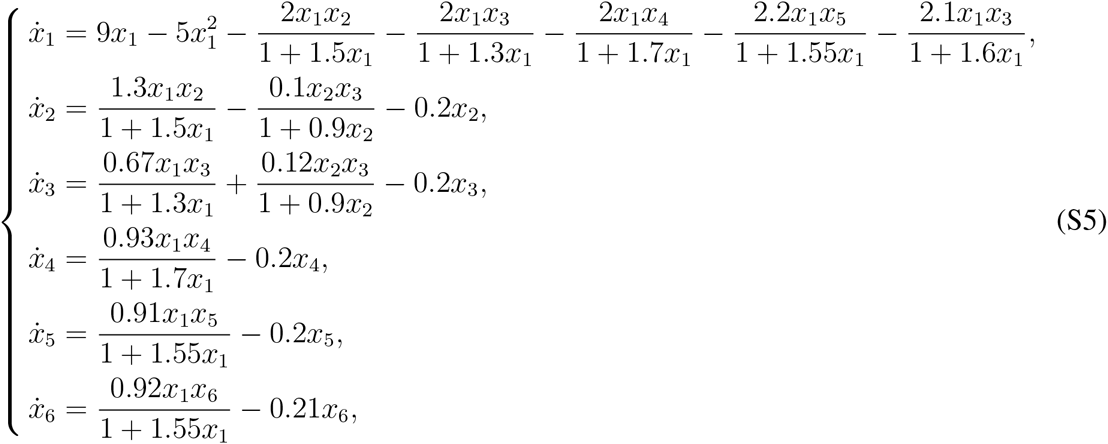

whose functional responses are of Holling Type II. The trajectories of the system and the interactions between species are shown in Fig. S3.

**FIGURE S3:**
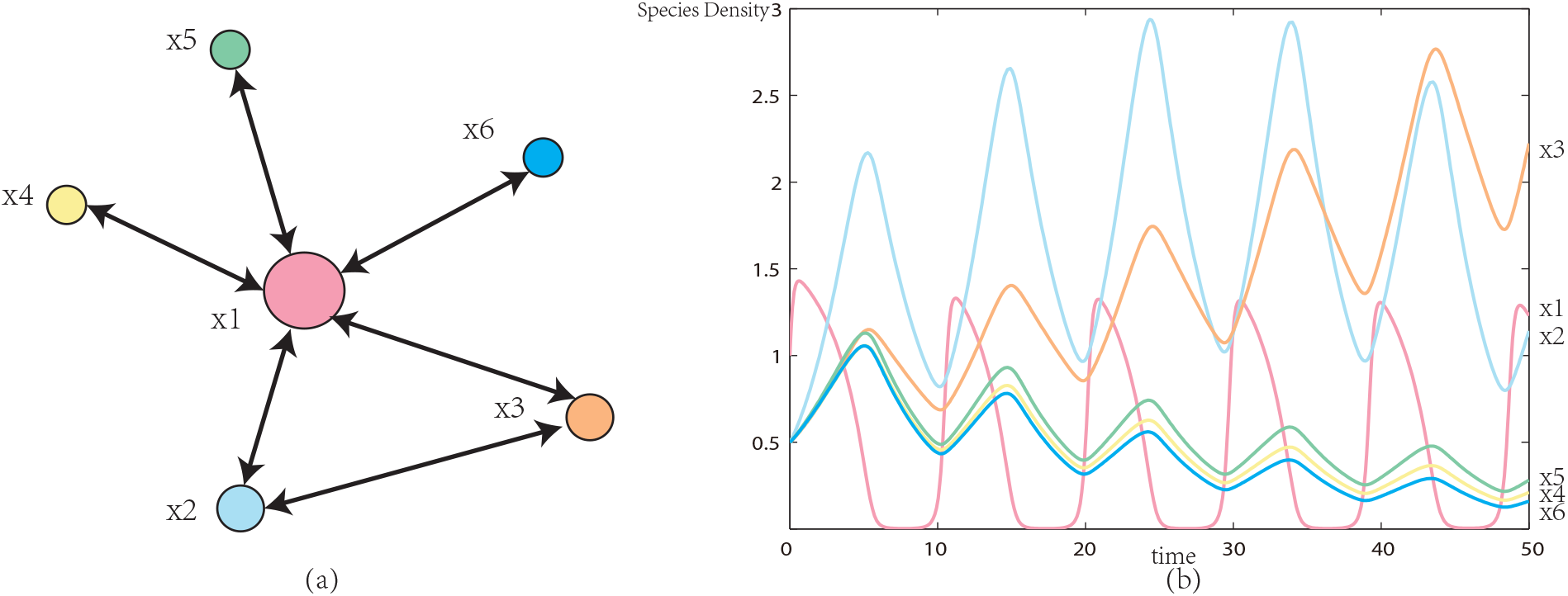
Synthesizing a six-species ecosystem. (a) With *x*_1_ playing as a central role in this synthetic system, pairwise interactions are constructed using Holling Type II functional responses. (b) Model parameters are tuned to obtain oscillations in all of the six species, providing informative time-series data for SR.

## 3. SR USING TEMPORAL DATA OF THE PREY IN ISOLATION

In the main text, we showed that additional temporal data of the prey in isolation 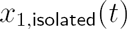 can be used to correctly infer complex functional responses. For this, we first estimate the derivative of this data 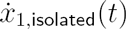 and use SR to find a good function 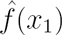 such that

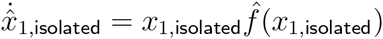

has a good fitness/complexity tradeoff. In the next step, we use the inferred 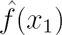 as prior information to the predator-prey model. Indeed, we can use the time-series data from the prey interacting with the predator {*x*_1_ (*t*), *x*_2_ (*t*)} to reverse-engineer the functional response 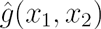 by using

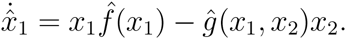

We found this approach very efficient, reducing the informativeness of the data needed to correctly infer *g*(*x*_1_, *x*_2_) and the parameters *m* and *µ.* In particular, this method allows us to correctly reverse-engineer a synthetic ecosystems with CM functional responses (results shown in Fig. 2b in main text).

Since it is natural to expect that the functional response of real ecosystems are at least as complex as the CM functional response, we applied the above method to the experimental data of Veilleux [18] with a predator-prey system of *P.aurelia* and *D.nasutum.* In such experiment, the authors reported data in which the isolated prey is cultured under the same conditions as the predator-prey system. We first interpolated these measurements using cubic splines and then sampled them every 0.1 days in order to generate the data 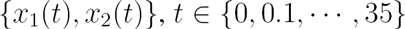. We also included the delay operator delay (*x*_*i*_(*t*)) = *x*_*i*_(*t* − 0.1) in the set of operators that the SR algorithm can use to build the candidate functions. Using this process, SR successfully inferred a biologically meaningful model with high fitness shown in Fig. S4.

**FIGURE S4:**
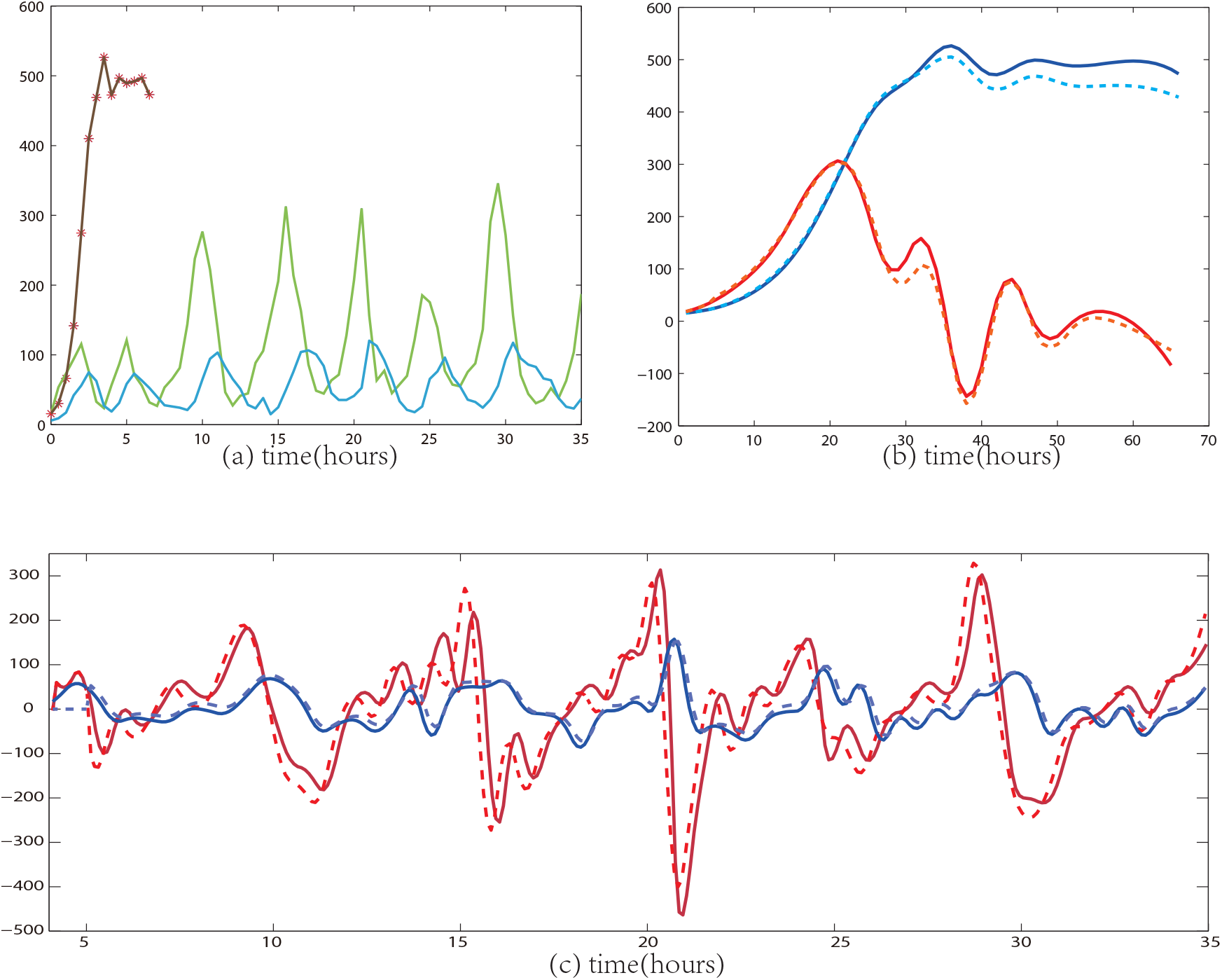
Inferring dynamics from experimental data of a predator-prey ecosystem. (a). We use splines to interpolate the original experimental data [18] of the prey in isolation (black), and the density of predator and prey (blue and green respectively). (b). we first use SR to discover the dynamics of prey in isolation, which accurately depicts both the prey density (blue) as well as prey variations (red), with dashed lines depicting the original datasets. (c). With the prior knowledge on the prey growth rate of prey, SR is able to discover meaningful models with 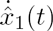 on the left hand side of the derived equation. We could recover both the prey density(blue) and prey variations (red) with the existence of predators, with dashed lines depicting original data in Fig. S4a.

## 4. USING A DICTIONARY OF POSSIBLE FUNCTIONAL RESPONSES

In case the temporal data is not informative enough, we can seed the SR algorithm with prior knowledge of possible functional responses. Indeed, we have explained that the main obstacle for directly using SR is that it cannot infer complex interactions with uninformative data, for instance, equilibrium points rather than limit cycles. Here we show that the prior knowledge about the system interactions —which is often available— can be used to decrease the necessary informativeness of the temporal data. This form of prior information can be regarded as the “dictionary” of possible structures of interactions revealing the temporal phenomenon, which we refer as the information blocks provided to the SR algorithm. We first collected and combined terms existing on the right-hand side of Equation. S3, such as 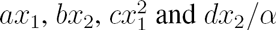 in Fig. S5. Note that for different types of functional responses, *α* denotes different structures of interactions in the denominator of *g*(*x*_1_, *x*_2_). We treat them as the prior knowledge we can get of interaction forms for inputs of SR in advance. Since it is rather difficult for SR to search through solution space to identify the exact structures of *α,* we move furthur to transform the equations, and put term *x*_1_*x*_2_ on the left side of the previous ODE, which is regarded as the output variable. Instead of providing SR algorithm with input species variables directly, we instructed SR with a set of possible terms existing in our interaction dictionary. To take the example of Holling type II functional response, on the right side now it should have terms 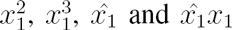. In this way the algorithm is provided with the previous knowledge for some extent of model structures hidden behind the temporal data. We listed all possible terms which may exist in the right hand side of the transformed target equations. In the next step, we could transform the reverse-engineering process of 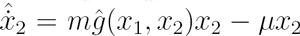 into a multi-gene SR problem of finding parameters for different units which are listed in our interaction dictionary. Some parameters simply equal to 0, indicating the absence of some types of functional responses. In this case, the multi-gene symbolic regression on the Matlab package GPTIPS2 [19] is efficient in extracting the meaningful blocks. Combined with a post-analysis on Pareto-front, it is possible to select the most insightful interaction units with the fitness/complexity tradeoff. Thus we can transform back the derived model into structure of Equation. S3, and we found it efficient in treating uninformative datasets.

**FIGURE S5:**
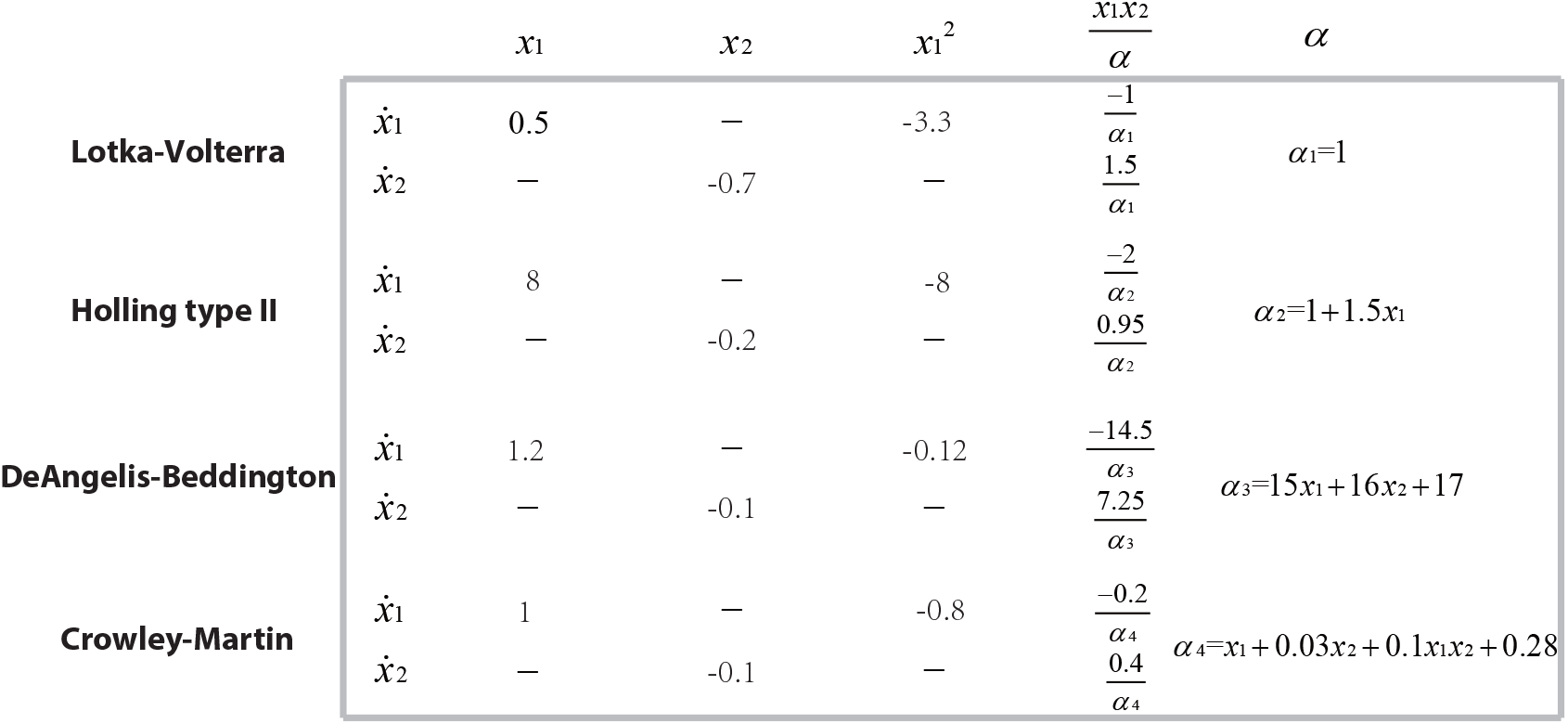
Model decomposition for multi-gene SR. When the data is not informative, which is shown in the case of Fig. S2, it is insightful to provide a dictionary of possible structures of interactions as prior knowledge. We firstly decompose right-hand side of original ODEs and transform the equations consisting of blocks of possible model structures.

Since this method decreases the needed informativeness of the data, it is also useful for inferring the dynamics of multi-species systems such as (S5). In the results shown in Fig. S3, at the initial stage, the *x*_4_, *x*_5_ and *x*_6_ satisfy the structure

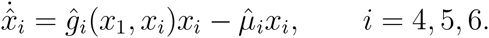

Therefore, applying the results of Section 2 we can infer 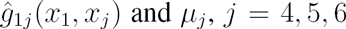, from informative time-series data {*x*_1_(*t*),…,*x*_6_(*t*)}. We then found the data was not informative enough for SR to recover the correct models for *x*_2_, *x*_3_. Hence we used the dictionary of possible functional responses together with multi-gene SR, allowing us to correctly infer 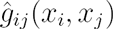 for *i* = 1, 2, 3 and *j* = 1,…,5. With all the recovered functional responses 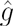 concerned with *x*_1_ provided as inputs, the SR algorithm was able to correctly infer the model of 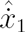, which has the highest complexity including pairwise interactions with all other 5 species. With the combination of multi-gene encoding of the model expressions and representative blocks transformation for the system dynamics, we manage to get rid of bloated equations or overfitting of specific models, and retrieve system dynamics with exact structures of function and accurate variable parameters.

